# The potyviral protein 6K1 reduces plant protease activity during *Turnip mosaic virus* infection

**DOI:** 10.1101/2021.09.05.459032

**Authors:** Sayanta Bera, Gabriella D. Arena, Swayamjit Ray, Sydney Flannigan, Clare L Casteel

**Affiliations:** Department of Plant-Microbe Biology and Plant Pathology, Cornell University, Ithaca, NY, 14850 USA; Laboratório de Biologia Molecular Aplicada, Instituto Biológico de São Paulo, São Paulo, SP, Brazil

**Keywords:** *Arabidopsis thaliana*, defence, jasmonic acid, MPMI, papain-like cysteine proteases, phytohormones, plant-microbe interactions, plant-virus interaction, protein turnover, transcriptomics, *Turnip mosaic virus*

## Abstract

- Potyviral genomes encode just 11 major proteins and multifunctionality is associated to most of these proteins at different stages of virus life cycle. The potyviral protein 6K1 is required for potyvirus replication at the early stages of viral infection and may mediate cell-to-cell movement at later stages.
- Our study demonstrates that the 6K1 protein from *Turnip mosaic virus* (TuMV) reduces the abundance of transcripts related to jasmonic acid biosynthesis and transcripts that encode cysteine protease inhibitors when expressed in *trans* in *Nicotiana benthamiana* relative to controls. Furthermore, 6K1 stability increases when lipoxygenase and cysteine protease activity is inhibited chemically, linking a mechanism to the rapid turnover of 6K1 when expressed in *trans*.
- Using transient expression, we show 6K1 is degraded rapidly at early time points in the infection process, whereas at later stages of infection protease activity is reduced and 6K1 becomes more stable, resulting in higher TuMV accumulation in systemic leaves. There was no impact of 6K1 transient expression on TuMV accumulation in local leaves.
- Together, these results suggest a novel function for the TuMV 6K1 protein which has not been reported previously and enhances our understanding of the complex interactions occurring between plants and potyviruses.

## INTRODUCTIONS

Viruses have evolved to perform all of the essential functions required to successfully infect a host, despite their very small genomes. At least two genetic strategies are observed that allow viruses to be more efficient with their limited coding potential (Elena *et al*., 2014). One of the strategies is to code for proteins from overlapping open reading frames (ORF) present in the viral genome (Schlub and Holmes, 2020). The overlapping ORFs strategy includes the presence of subgenomic RNA and a frameshift of the starting codon of an ORF (Schlub and Holmes, 2020). Another strategy is the multifunctionality of the viral encoded proteins (Callaway *et al*., 2001; Valli *et al*., 2018). Some viral proteins are known to perform multiple critical functions in the virus life cycle, such as genome replication, encapsidation, intercellular movement, long-distance movement, RNA silencing suppression, or vector transmission (Deng *et al*., 2015; May *et al*., 2020; Valli *et al*., 2018). These above-mentioned strategies are not mutually exclusive and help viruses to circumvent the problem of small genome size.

The multifunctionality of viral proteins can be regulated spatially and temporally, and is dependent on ecological conditions and stages of the viral life cycle. A study on the viral protein nuclear inclusion a protease (NIaPro) demonstrated that apart from the proteolytic activity, it also has a role in plant-aphid interactions by re-localizing outside of the nucleus of a plant cell when the aphid vector is present (Bak *et al*., 2017). Thus, Bak *et al*., (2017) demonstrates the role of ecological interactions in modulating the location and function of potyviral proteins. Post-translational modification of viral proteins can also be used to change their physio-chemical properties, such as stability. For example, the coat protein (CP) of *Potato virus A* is degraded rapidly at early time points in the infection process, whereas at later stages the CP becomes more stable, when systemic infection and encapsidation are critical (Ivanov *et al*., 2003). It was determined that phosphorylation of the CP was responsible for the dynamic stability of CP, correlating with the viral life cycle (Ivanov *et al*., 2003). Another way to regulate viral protein stability is by interfering with the host’s degradation machinery. For example, proteins encoded from the genomes of *Cucumber mosaic virus* (CMV), *Cauliflower mosaic virus* (CaMV), *Barley stripe mosaic virus* (BSMV), *Tomato yellow leaf curl virus* (TYLCV), *Cotton leaf curl Multan virus* (CLCuMuV), and *Turnip mosaic virus* (TuMV) are known to interact with proteins in the autophagy or the ubiquitin-proteasome degradation pathway, affecting viral protein turnover (Cheng and Wang, 2017; Hafrén *et al*., 2017; Hafrén *et al*., 2018; Ismayil *et al*., 2019; Li *et al*., 2018; Yang *et al*., 2018).

Potyviruses have single-stranded positive-sense RNA genomes which characteristically contain one ORF. The single ORF encodes a large polyprotein that is cleaved into 10 functional proteins by virus-encoded proteases (García *et al*., 1992; Revers and García, 2015). Some potyviruses contain another small overlapping ORF which codes the PIPO protein and is located within the P3 cistron of the polyprotein (Chung *et al*., 2008). While each potyviral proteins is known to perform multiple functions, not much is known on the function of the 6K1 protein, a small 6 kDa protein. Early research suggested 6K1 may play a role in viral replication and in cell-to-cell movement (Hong *et al*., 2007; Johansen *et al*., 2001), however research related to 6K1 has been limited over the past two decades due to its small size, instability due to high protein turnover, and low protein expression levels (Cui and Wang, 2016; Geng *et al*., 2017; Kekarainen *et al*., 2002; Merits *et al*., 2002; Waltermann and Maiss, 2006). Despite these challenges, Cui and Wang, (2016) presented evidence that 6K1 is required during the early stages of *Plum pox virus* (PPV) replication and recently a study demonstrated that the 6K2 protein, which is considered as a marker for viral replication complex (VRC), recruits 6K1 to the potyvirus replication complex (Geng *et al*., 2017), confirming earlier findings. Surprisingly, the 6K1 protein, which also contains a hydrophobic transmembrane domain like the 6K2 protein, is found in the soluble protein fraction, while 6K2 is found in membrane fraction (Cui and Wang, 2016; Jiang *et al*., 2015).

Although using a mutated infectious clone simulates the natural viral infection process, in the case of 6K1 mutations are lethal for virus survival (Cui and Wang, 2016; Kekarainen *et al*., 2002; Merits *et al*., 2002). The goal of this study was to further our understanding of 6K1’s function using a model system consisting of the potyvirus TuMV and *Nicotiana benthamiana* as a host. To circumvent the difficulty of no or reduced viral infection from mutating 6K1 in an infectious clone, we assayed the function of 6K1 by ectopically expressing it in uninfected plants and in the presence of a wildtype TuMV infection. This strategy allowed us to investigate in detail additional functions associated with the 6K1 protein during the virus life cycle and circumvent issues with 6K1 protein stability. Our results suggest jasmonic acid and papain-like cysteine proteases may play a role in changes in 6K1 stability and function during the infection process. Understanding the molecular mechanism behind the 6K1 protein degradation will also pave the way for future studies on the critical multifunctionality associated with viral proteins and successful viral infection in plants.

## RESULTS

### The ectopically expressed 6K1 protein is degraded by cysteine proteases

We first examined stability of 6K1 using our construct, transient expression, and fluorescent microscopy at 72 hours post agro-infiltration (hpi). Consistent with previous reports (Cui and Wang, 2016), 6K1:GFP fluorescence was only weakly visible compared to GFP controls using confocal microscopy (Fig. 1A). The kinetics of GFP and 6K1:GFP accumulation were monitored by western blot analysis using antibodies that recognize GFP (Fig. 1B). For GFP, protein expression was stable until 72 hpi, whereas, for 6K1:GFP, protein expression was only detected at 48 hpi (Fig. 1B). Next, chemical inhibitors were used that target the different protein degradation pathways: 3MA (autophagy inhibitor), MG132 (cysteine protease and proteasomal inhibitor) and E64 (papain-like cysteine proteases inhibitor) (Fig. 1C). Although 3MA and MG132 did not impact 6K1:GFP protein accumulation relative to the GFP control, E64 increased the 6K1:GFP protein accumulation significantly relative to the control (Fig.C). Next, to assay if 6K1 protein stability is also affected post transcriptionally, an experiment was performed with a viral suppressor of RNA silencing (VSR), P19, co-infiltrated with GFP/6K1:GFP. Western blot analysis revealed that 6K1:GFP accumulation was higher in the presence of P19 in comparison to the control (Fig. 1C). Considering all the results, 6K1 protein stability in uninfected plant is regulated at both the protein synthesis and degradation level.

**Fig. 1.**
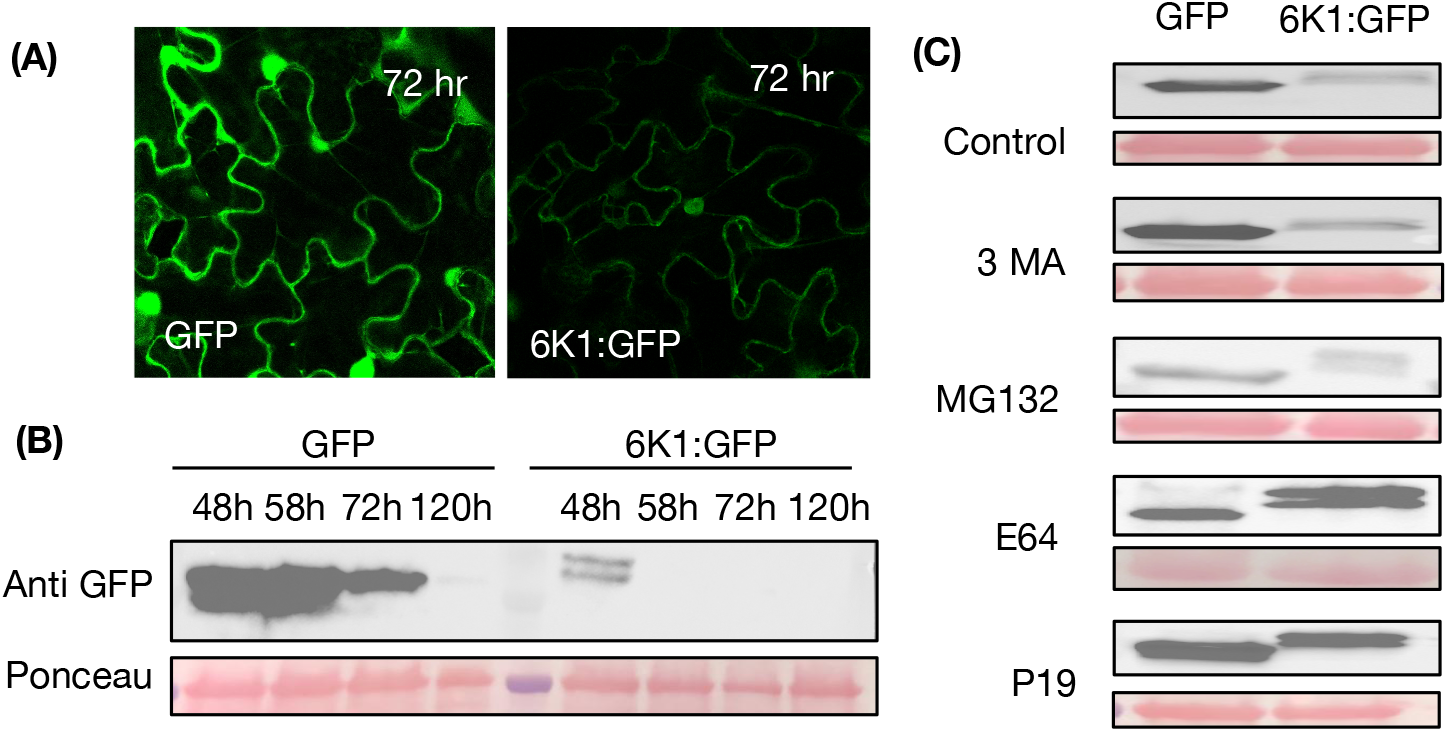
The ectopically expressed 6K1 protein is degraded by cysteine proteases. The construct GFP and 6K1:GFP were agroinfiltrated in *N. benthamiana* leaves. (A) Pictures were taken using a confocal laser scanning microscope with a 40X objective (images are a single section of 1-μm thickness). The green fluorescence indicates the protein accumulation of GFP and 6K1:GFP 72 hr post agroinfiltration in a *Nicotiana benthamiana* leaf. (B) Western blots were performed with proteins extracted from the agroinfiltrated leaves collected overtime. (C) Western blot analysis of transiently expressed GFP and 6K1:GFP was assessed in the presence of chemical inhibitors of the autophagy protein degradation pathway (3MA), the proteasome protein degradation pathway (MG132), cysteine proteases (E64), and P19, an RNA interference silencing suppressor. Constructs were co-agroinfiltrated, while chemical inhibitors were infiltrated 48 hours after agroinfiltrations. For each sample, an equal volume was loaded into each well of an SDS-PAGE gel. Anti-GFP was used in both western blots and Ponceau staining was performed to check for loading control. All western blots are representative of at least two replicates which contained 3 plants per treatment (N=3).

Protease inhibitors contribute to protein regulation in plants by preventing protein turnover during development and senescence in plants (Habib and Fazili, 2007). We hypothesized protease inhibitors may also be reduced, which may contribute to the increased 6K1 protein turnover when expressed in trans shown here and in previous reports (Cui and Wang, 2016). To determine if protease inhibitors are decreased in plants expressing 6K1, we measured transcript abundance of the *N. benthamiana Cystatin* protease inhibitor (Niben101Scf00862g02050.1) with the highest identify to a tomato cystatin that was previously implicated in plant-insect interactions (Goulet et al., 2008). Relative to the control treatment (GFP), *Cystatin* transcripts were significantly reduced in presence of 6K1 (Fig. 2A). To determine if other post-translational regulation pathways contribute to 6K1 turnover, we next measured the transcript abundance of markers for different protein degradation pathways (*NBR1* for autophagy, *RPN11* for proteasome) in the same treatments. *NBR1* and *RPN11* expression were both highly induced in the presence of 6K1 protein relative to the control (Fig. 2B, C), suggesting multiple pathways contribute to turnover.

**Fig. 2.**
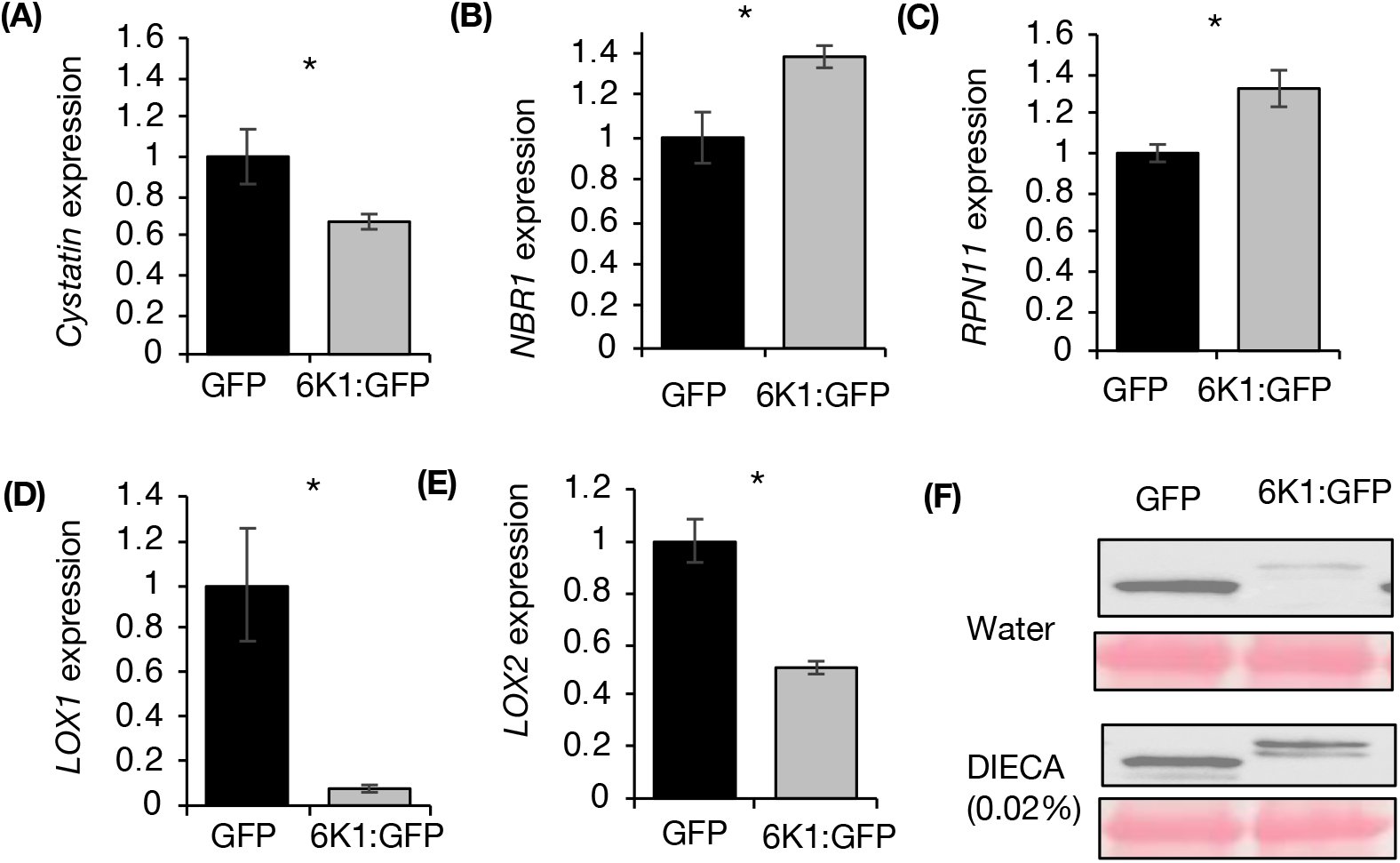
6K1 expression inhibits transcripts related to protease inhibitors and jasmonic acid accumulation. RNA was extracted from *Nicotiana benthamiana* leaves transiently expressing the GFP or 6K1:GFP. Transcript abundance of markers for different protein degradation pathways were measured using qRT-PCR: (A) *Cystatin*, a protease inhibitor, (B) *NBR1*, related to autophagy, and (C) *RPN11*, related to proteasome degradation. We next measured the transcript abundance of (D) *LOX1* and *(E) LOX2*, related to jasmonic acid biosynthesis. The relative quantification was performed by using *actin* as reference gene and GFP treatment as the calibrator. Each result is the mean from 5 replicated plants ± SE. The stars denotes if the mean differences were significantly different at *P* < 0.05 as determined from either *t*-test or, Kruskal-Wallis test. (F) The GFP or 6K1:GFP constructs were agroinfiltrated in *N. benthamiana* leaves and a lipoxygenase inhibitor (DIECA) was sprayed on half the plants 48 hr later. At 60 hr post-agroinoculation proteins were extracted and SDS-PAGE gels were run with equal volume (9μl) of each samples. Anti-GFP was used in both western blots and Ponceau staining was performed to check for loading control. The western blot is representative of at least two replicates which contained 3 plants per treatment (N=3).

### 6K1 expression inhibits transcripts related to jasmonic acid accumulation

We hypothesized that 6K1 might also alter jasmonic acid accumulation in uninfected host plants because jasmonic acid is known to regulate the production of protease inhibitors (Farmer *et al*., 1992). To address this, we measured the abundance of two transcripts related to JA biosynthesis, *LIPOXYGENASE1* (*LOX1)* and *LIPOXYGENASE2 (LOX2*), in *Nicotiana benthamiana* leaves transiently expressing the GFP or 6K1:GFP protein. 6K1:GFP expression significantly inhibited *LOX1, LOX2 transcripts* and JA accumulation compared to controls (Fig. 2D,E). To determine if inhibition of LOX1 and LOX2 is relevant to 6K1:GFP stability, plants that were previously agro-infiltrated with either GFP/6K1:GFP, were sprayed with an lipoxygenase inhibitor, DIECA (Farmer et al. 1994), 48 hours after infiltration (Fig. 2F). Western blot analysis revealed that 6K1:GFP accumulation was higher in the presence of DIECA in comparison to the GFP control (Fig. 2F).

### Transcriptome wide analyses revealed that aphid and TuMV differentially affect host protein degradation pathways in *A. thaliana*

Next we performing RNA-seq to examine the transcriptome of *A. thaliana* in following different treatments: infected with TuMV, infested with aphids or, when both TuMV and aphids were present. Differential gene expression analysis revealed TuMV infection had a greater impact on transcriptional changes compared to aphid infestation (Fig. S1A, B, C). Overall, 188 and 368 genes were differentially expressed exclusively in response to aphid and TuMV treatments, respectively, whilst 19 genes were regulated in both treatments (Fig. S1B). Only 15 transcripts were shared among all treatments, and the greatest number of transcripts were regulated in the treatment with both aphids and TuMV compared to controls (Fig. S1C). Gene set enrichment analysis (GSEA) was used next to determine which biological processes were over-represented in each treatment (Table S1 and S2). For aphid treatment, categories related to responses to herbivore, JA, ethylene, and abscisic acid along with JA and ethylene biosynthetic processes were enriched (Table S2), which are in line to our previous published data (Bak *et al*., 2019; Casteel *et al*., 2015; Hillwig *et al*., 2016). For TuMV treatment, GSEA analysis indicated biological process related to salicylic acid biosynthesis and responses to JA and ethylene were significantly enriched (Table S1), which were also consistent to our previous findings (Bak *et al*., 2019; Casteel *et al*., 2015). Although we observed that though TuMV infection caused the most substantial changes in gene expression (Fig. S1), it was connected to fewer biological processes (130) compared to aphids (188). Nevertheless, highest number of biological processes were found to be regulated when both TuMV and aphids were present (230, Table S3).

To study the impact of treatments on specific protein degradation pathways (autophagy, proteasome, and protease inhibitors/proteases), we next searched the transcriptome for transcripts whose levels changed >1.5 times (upregulated or downregulated; *P* < 0.1) relative to the mock in either of the treatments (Fig. 3). Our data shows TuMV, aphids, or both treatments regulated 5 genes related to autophagy, 10 genes related to the proteasome, 14 related to protease inhibitors, and 46 related to proteases (Fig. 5A,B,C,D). Specifically, TuMV significantly induced the expression of *NBR1* (AT4G24690), which has been shown to be up-regulated previously by TuMV and to have a pro-viral function (Hafrén *et al*., 2018). The autophagy and proteasome pathway was found to be most differentially regulated when either TuMV or both TuMV and aphids were present and least when aphids were present alone. Aphids alone and aphids with TuMV had the greatest impact on protease inhibitor genes among all the treatments (Fig. 3C). The greatest number of genes related to proteases were significantly regulated when TuMV was present either alone or with aphids, whereas only aphids did not have as large of an impact on proteases (Fig. 3C). Taken together these results suggest TuMV infection downregulated mostly protease genes and upregulated some authophagy and proteasome related genes, while aphid feeding was mostly associated with the downregulation of protease inhibitor genes (Fig. 3A,B,C,D).

**Fig. 3.**
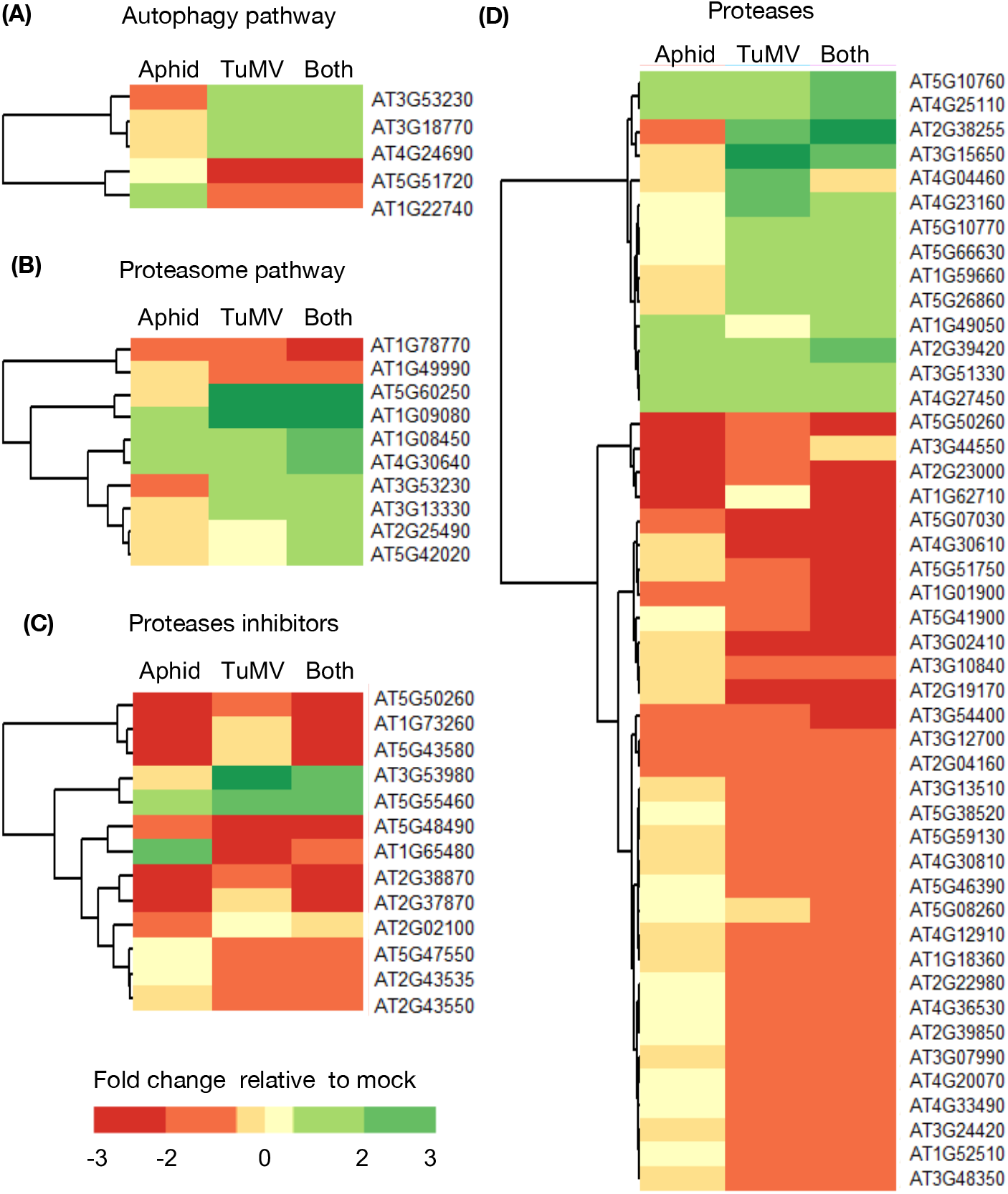
TuMV infection and aphid feeding significantly impact protein degradation pathways in *Arabidopsis thaliana*. RNA-seq was performed with mock-inoculated *A. thaliana, A. thaliana* one week after TuMV infection (TuMV), *A. thaliana* 48 hr after infestation with the *Myzus persicae* aphid (Aphid), a vector of TuMV, or from plants with both treatments (Both). Each sample represented a pool of two plants and three samples were taken per treatment (N = 3, 6 plants total). Heatmaps show genes that where at least 1.5 times differentially expressed relative to mock (*P*-value < 0.1). Differentially expressed genes were grouped according to the following protein degradation pathways: Autophagy (A), Proteasome (B), Protease inhibitors (C) and, Proteases (D).

### 6K1 protein stability increases and protease activity decreases during TuMV infection of *N. benthamiana*

Next, we expressed 6K1:GFP with and without an infectious clone of TuMV in *Nicotiana benthamiana*. The kinetics of 6K1:GFP protein accumulation were monitored by western blot analysis using anti-bodies that recognize GFP and with UV light. Without TuMV infection, a band was detected at 48 and 58 hpi and had an estimated molecular weight of 33kD, which is around the expected size of the 6K1:GFP fusion (Fig. 4A). After 58 hpi, 6K1:GFP expressed alone was not detected in the western blot analysis (Fig. 4A) or by UV light (Fig. S4). In contrast, 6K1:GFP in the presence of TuMV was detected at all time points in western blot analysis (Fig. 4A) and was visible with UV light in plants 120 hpi (Fig. S4). At early time points (48 and 58hpi), protein levels of 6K1:GFP were reduced in the presence of TuMV compared to without the infectious clone (Fig. 4A). As a control we conducted a similar experiment using GFP with and without TuMV (Fig. 4B) and found similar results (Fig. 4A). As the previous experiments demonstrated protease inhibitors and proteases are critical for 6K1 turnover (Fig. 2), biochemical assays were performed to quantify the total protease activity in the same treatments at 120h. Total protease activity was reduced when 6K1:GFP was expressed in the presence of TuMV compared to without TuMV (Fig. 4C), however there was no difference between the GFP treatments (Fig. 4D).

**Fig. 4.**
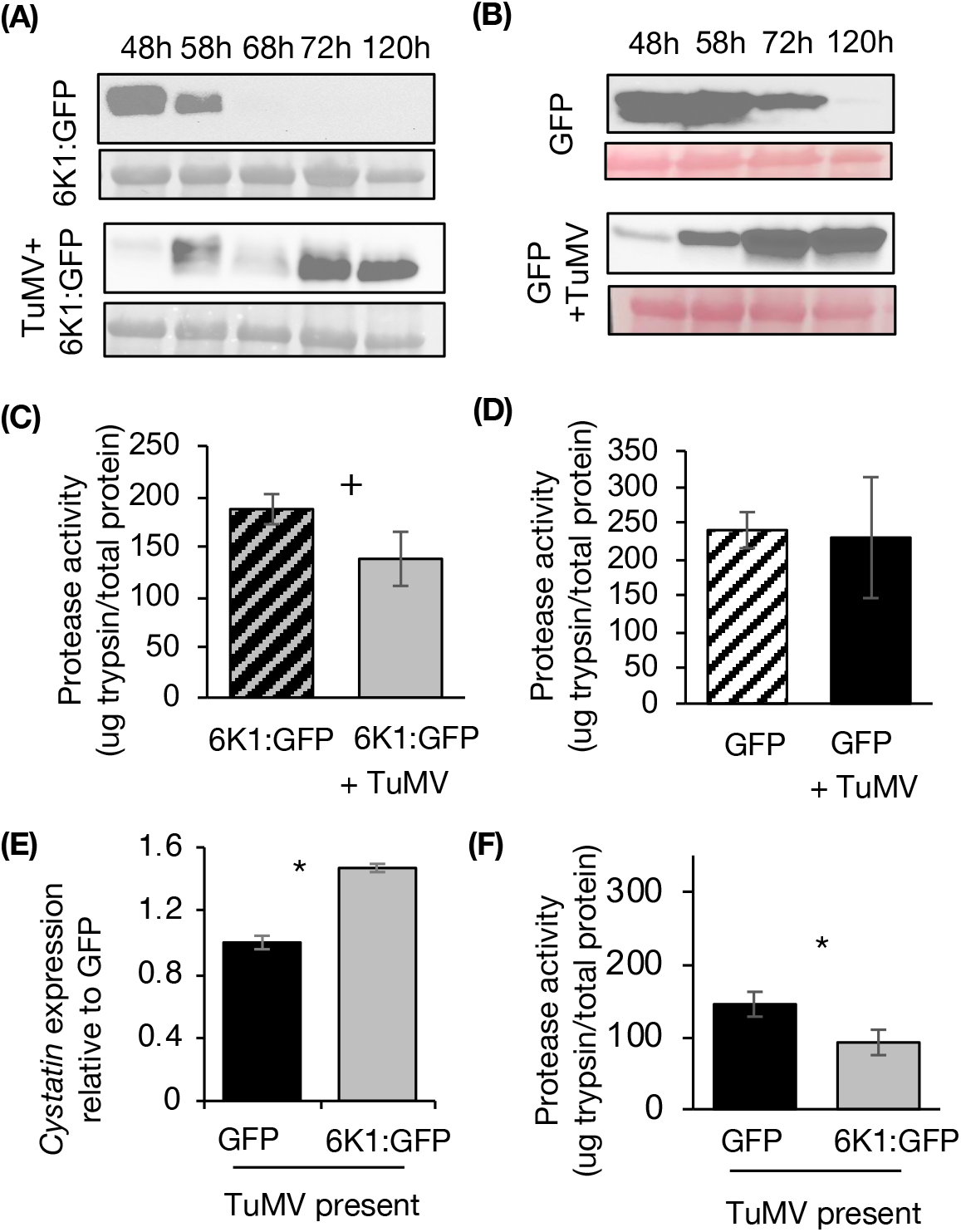
6K1:GFP protein stability increases and protease activity decreases during TuMV infection. Western blots were performed with *Nicotiana benthamiana* leaves expressing (A) 6K1:GFP or (B) GFP with and without TuMV over time. For each sample, an equal volume (10μl) was loaded into each well of an SDS-PAGE gel. Anti-GFP was used in both western blots and Ponceau staining was performed as to check for loading control. *N. benthamiana* leaves were agro-inoculated with TuMV and 5 days later (C) 6K1:GFP or (D) GFP constructs were agro-inoculated into the same leaves or into healthy control plants. Tissue was collected 60 hr later and protease activity was quantified. (E) Relative expression of *Cystatin* transcripts and (F) total protease activity were quantified in *N. benthamiana* leaves co-agro-inoculated with the GFP or 6K1:GFP protein and TuMV 60hr post agroinfiltration. Each result is mean from five biological replicates in (B), from 6-12 biological replicates in (C,D,EF) and a t-test was used in (B,C,D,E,F) to check for significance at a *P-value* of * < 0.05 and + < 0.1. All western blots are representative of at least two replicates which contained 3 plants per treatment (N=3).

To further investigate the role of 6K1 in inhibiting protease activity during TuMV infection, we next measured the abundance of the *Cystatin* transcript and total protease activity in plants transiently expressing GFP or 6K1:GFP in the presence of TuMV (Fig. 4E,F). In this experiment, transient expression of the 6K1 protein significantly induced transcript accumulation of the protease inhibitor, *Cystatin*, compared to the GFP control in the presence of TuMV (Fig. 4E). There was also a highly significant inhibition of protease activity when 6K1 was expressed in the presence of TuMV compared to the GFP control (Fig. 4F). These results suggest 6K1 expression can decrease plant protease activity, however as 6K1:GFP was not able to inhibit *Cystatin* in uninfected plants (Fig. 1), additional TuMV proteins or TuMV-induced plant proteins are required for this function during infection.

### TuMV accumulation is increased in systemic leaves in the presence of the ectopically expressed 6K1:GFP

Our results above demonstrate 6K1 degrades rapidly at earlier time points in infection, whereas at later stages of infection it becomes more stable, when systemic infection is more important. To determine the impact of transiently expressed 6K1 on virus movement, leaves were infiltrated with either TuMV or TuMV co-infiltrated with GFP or 6K1:GFP and the abundance of coat protein transcripts were quantified in local and systemic leaves. Virus coat protein transcripts accumulated to a similar level in local leaves in which only TuMV was infiltrated compared to leaves co-infiltrated with TuMV and either GFP or 6K1:GFP (Fig. 5A). In systemic leaves greater amounts of TuMV CP transcripts were detected when 6K1:GFP was co-infiltrated with TuMV relative to the control treatment (Fig. 5B).

**Fig. 5.**
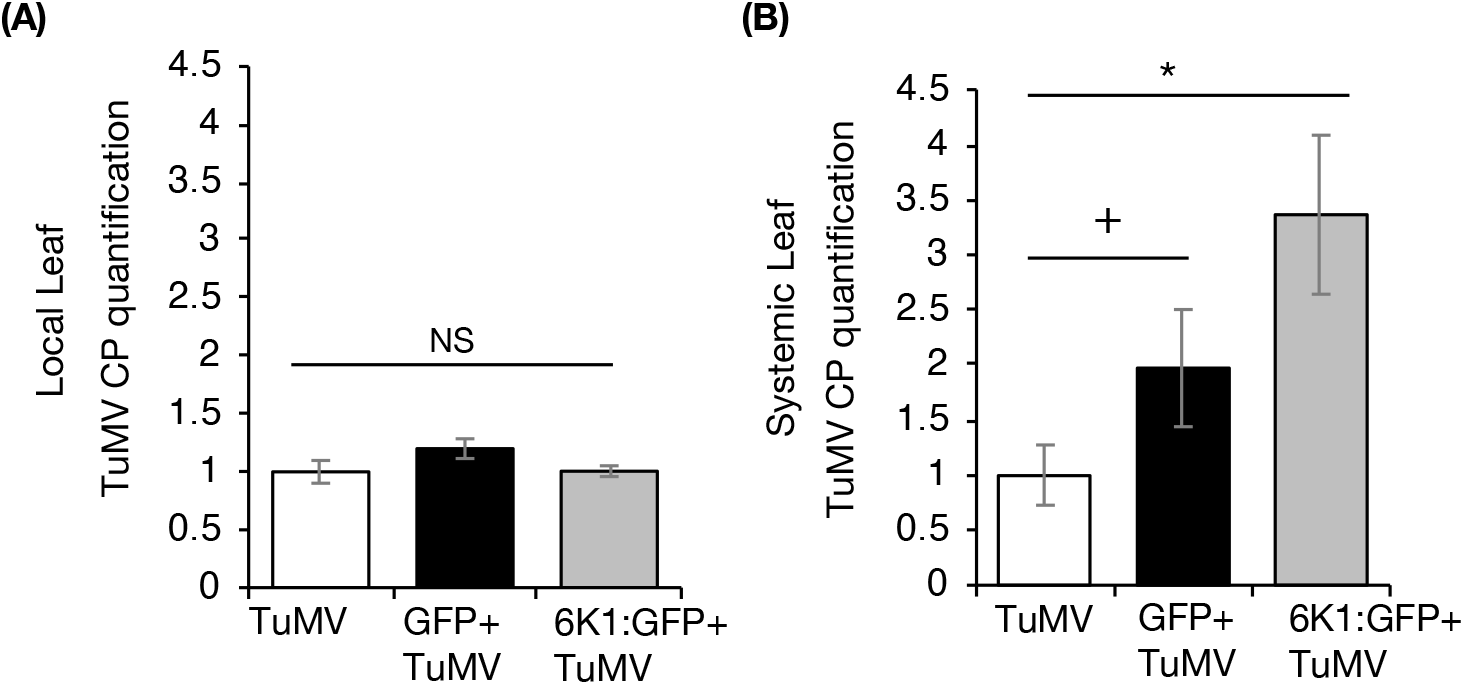
TuMV accumulation is increased in systemic leaves in the presence of the ectopically expressed 6K1:GFP. GFP or, 6K1:GFP constructs were co-infiltrated with TuMV into *N. benthamiana* leaves. In control treatment, only TuMV was agro-infiltrated. At 60 hr post infiltrations, coat protein specific primers were used for quantification of viral RNA relative to the actin in the local and systemic leaves. Significance was determined using differences at a *P-value* of * < 0.05 and + < 0.1 as determined from a *least significance difference* (LSD) test (*n* = 5, mean ± SE).

## DISCUSSIONS

In this study, we demonstrate that the stability of the 6K1 protein is dynamic and increases over time in the presence of TuMV (Fig. 4). Decreased stability of the ectopically expressed 6K1 was due to post-transcriptional changes and due to changes in protein degradation (Fig. 1). It was previously reported that the expression of 6K1 protein *in-vivo* is quite low and affinity purification was required to detect PPV’s 6K1 protein during viral infection (Waltermann and Maiss, 2006). In our study transient expression of the 6K1 protein was lowest 48 hpi relative to other time point in virus infected plants and increased overtime (Fig. 4A). A similar observation was reported in Waltermann and Maiss (2006) where the 6K1 protein was not detected at 48 hpi in virus infected plants, but at 96 hpi they were able to detect it. It is important to note when a second GFP-tagged copy of PPV’s 6K1 was expressed in *cis* from an infectious clone, increased stability of 6K1 was not observed overtime (Cui and Wang, 2016). In our construct the cleavage site was retained at the C-terminus of 6K1 before the GFP sequence, so that during virus infection 6K1 could still be cleaved from GFP to avoid any negative impacts on function. It is tempting to speculate that early in the potyvirus infection process the 6K1 protein may be inhibited by the virus or plant, while later in the infection process 6K1 stability is increased enabling new functions. For example, we demonstrated ectopic expression of 6K1 increased viral accumulation in systemic leaves (Fig. 5), suggesting 6K1’s increased stability may play a role in regulating viral movement.

We observed higher accumulation of the transiently expressed 6K1 protein in the presence of P19 VSR, which suggests the 6K1 mRNA is unstable and a target of host RNA silencing machinery (Fig. 1C). As the activation of host RNA silencing machinery is one of the primary defence responses that target viruses (Calil and Fontes, 2017; Wang *et al*., 2012), it seems reasonable that parts of the viral RNA genome will be potent activators of RNA silencing. Nevertheless, potyviruses code for VSRs (HCPro and VPg) to ensure successful virus infection (Eskelin *et al*., 2011; Valli *et al*., 2018). Indeed, the work of LaTourrette and Garcia-Ruiz., (2021) showed that HC-Pro is mostly bounds to 21 nt siRNAs of viral origin thus, preventing host gene silencing pathway from targeting viral RNA. Recently, papain-like cysteine proteases (PLCPs) have been showed to play an important role in host defences against many pathogens including viruses (Bar-Ziv *et al*., 2012; Bar-Ziv *et al*., 2015; Misas-Villamil *et al*., 2016). We show here proteases do have a role in degradation of the ectopically expressed 6K1 protein, as protein accumulation increased in the E64 treatment, a chemical inhibitor of PLCPs (Fig.1C). Many PLCPs are localised in autolysosomes, suggesting a role of the autophagy pathway in the degradation of 6K1 protein (Bárány *et al*., 2018; Nakahara *et al*., 2012) However, there was no change in 6K1 protein accumulation in the treatment with 3-MA, an inhibitor of the autophagy pathway (Fig. 1C). Surprisingly, we did observe 6K1 expression causes a significant increase in *NBR1* and *RPN11* (Fig 2B,C), two transcripts related to autophagy and the proteasome degradation pathway respectively, which suggest additional studies are required to determine their role in 6K1 stability. We go further and demonstrate that protease activity, and papain-like cysteine protease inhibitors are regulated by 6K1 (Fig. 4) and TuMV infection (Fig. 3). Overall, our data suggests that the protein stability of ectopically expressed 6K1 is regulated at multiple levels and paves the way for additional investigations on the 6K1 protein and the complex molecular mechanisms associated with a successful virus infection.

Previous studies have shown that 6K1 has a role in viral replication and may mediate cell-to-cell movement (Cui and Wang, 2016; Geng *et al*., 2017; Hong *et al*., 2007; Lõhmus *et al*., 2016). Our data provides evidence of a novel function associated to 6K1 protein *i*.*e*., inhibition of JA biosynthesis transcripts (Fig. 2D,E,F). It is well established that phytohormones mediate many different components of plant-virus-insect interactions and may regulate virus transmission (Farmer et al., 1992; Abe *et al*., 2012; Bera *et al*., 2020; Lewsey *et al*., 2010; Tian *et al*., 2015; Parizad and Bera 2021; Basu *et al*., 2021; Lee *et al*., 2021). Inhibition of JA in the presence of 6K1 protein may indicate a possible role of 6K1 in mediating ecological interactions. Indeed, it was shown that 6K1 decreases aphid fecundity (Casteel *et al*., 2014) and aphids induce JA in plants (Casteel et al., 2015), suggesting a positive effect of JA on aphids. Taken together, it can be speculated that 6K1-mediated inhibition of JA may result in the poor performance of aphids we previously observed.

As viral proteins are often associated with more than one function thus, it is critical to assay their function throughout virus life cycle and under different ecological conditions. Our results also support earlier observation of 6K1’s role in systemic movement using ectopic expression with and without the virus instead of mutating infectious clone (Fig. 5). As viral RNA acts as an open reading frame and may also form a functional RNA element, mutating a part of the viral genome can be detrimental to virus survival (Ilyas *et al*., 2021; Liu *et al*., 2021). The evolution of potyviruses with plants has progressed for about 15,000 to 30,000 years and the emergence of new potyviral species are still being documented (Gibbs *et al*., 2020; Parizad *et al*., 2018). Further, potyviruses are among the most widely distributed pathogens in crops, hampering the production and the quality of food (Karasev and Gray, 2013; Moratalla-lópez *et al*., 2021). While our study paves the way for more thorough investigation of the 6K1 protein, its’ multifunctionality, and role in plant-virus-aphid interactions, a thorough understanding of the underlying molecular mechanism causing potyviruses disease will be required to develop innovative intervention strategies and prevent viral epidemics in the future.

## EXPERIMENTAL PROCEDURES

### Plants and growth conditions

*Nicotiana benthamiana* and *Arabidopsis thaliana* plants were grown in growth chambers under controlled conditions (25/20 °C day/night with a photoperiod of 14/10 h day/night) at a relative humidity of 50% and a light intensity of 200 mmol m^−2^ s^−1^. Plants were grown for 3 to 4 weeks and were used in experiments before flowering, unless otherwise noted.

### Virus inoculation

TuMV was propagated from the infectious clone pCAMBIA:TuMV, kindly provided by Prof. Jean-Francois Laliberte. In all the experiments using pCAMBIA:TuMV, virus inoculation was mediated by the agro-infiltration of the agrobacterial culture after diluting to an optical density of 0.03 at 600 nm, unless otherwise noted.

### Plasmid constructs

The 6K1:GFP constructs and its derivatives were produced using the Gibson cloning kit (New England Biolabs, Ipswich, MA) following the manufacturer’s instructions. Briefly, compatible gene-specific Gibson primers were designed to perform PCR and the PCR product and the digested pMDC32 vector were joined using Gibson assembly. To clone 6K1 and GFP into pMDC32 plasmid, p35:TuMV/GFP (Casteel *et al*., 2014) was used as a template to amplify 6K1 and GFP separately. pMDC32:6K1, pMDC32:6K1:GFP, and pMDC32:GFP were then assembled as above using the Gibson kit.

### Transient protein expression in *Nicotiana benthamiana*

All the plasmid constructs were introduced into *Agrobacterium tumefaciens* GV3101 separately by heat shock and selected on LB plus 10 μg ml^-1^ of rifampicin and 50 μg ml^-1^ of kanamycin. One fresh colony was selected and grown overnight in liquid culture with the same antibiotic selection as before. The pellet of the culture was resuspended in 10mM MgCl_2_ and 150 μM acetosyringone and left at room temperature for 2 – 3 h in a dark room. The solution containing the agrobacterial culture was then diluted to an optical density of 0.2 at 600nm for transient expression experiments and at 0.1 for co-infiltrating with the TuMV infectious clone. Single leaves from 4-week-old *N. benthamiana* plants were then agro-infiltrated with the agrobacterium culture suspended in MgCl_2_ and acetosyringone. After agro-infiltration, leaf tissue (100 mg) was collected 48 hours post infiltration and, thereafter samples were taken at 72 hours, 5 days, from separate plants and according to the individual experiment’s design. Expression was verified by microscopy, RT-PCR and/or western blot analysis as described below.

### Chemical inhibitors treatments

To investigate how the ectopically expressed 6K1 protein gets degraded, several assays with chemicals were performed that inhibit specific pathways of protein degradation in plants. MG132 (Sigma-Aldrich, St. Louis, MO), and 3Methyladenine (3MA) (Tci America, Tokyo, Japan) were used to inhibit the proteasomal degradation and autophagy pathways, respectively and, E64 (Sigma-Aldrich, St. Louis, MO) was used to inhibit the cysteine proteases. The 3MA (10mM) solution was prepared by dissolving it in phosphate buffered saline (PBS) containing 2% DMSO. MG132 (50μM) was prepared in PBS and E64 (50 and 100μM) was prepared in water. Forty-eight hours after 6K1 agro-infiltration, chemical inhibitors (1 ml) were infiltrated in the previously agro-infiltrated leaves. Post agro-infiltration samples were collected at 60hpi.

### Microscopy

Leaves were cut into small pieces and placed on glass slides and observed with a Leica TCS SP5 confocal laser scanning microscope system (Leica Micro-systems, Bannockburn, IL, USA). GFP fluorescence was detected with excitation at 488nm and emission capture at 500–530 nm. Images were captured at 2,361 mm intervals using a 10x objective.

### Western blotting

Leaf tissue was collected and flash frozen in liquid nitrogen. Later it was crushed in a lysis buffer (10mM sodium citrate, 1% SDS, 30mM NaCl, 0.4% 2-mercaptoethanol, 2X EDTA-free protease inhibitor cocktail) and boiled for 10 min in a 1.5 ml tube. The supernatant was mixed with an equal volume of loading dye and fractionated by a 12% SDS–polyacrylamide gel electrophoresis gel under reducing conditions. Afterwards protein bands were transferred to a nitrocellulose membrane using a transfer apparatus according to the manufacturer’s protocols (Bio-Rad, Hercules, CA). After incubation with 5% nonfat milk in TBST (50mM Tris-Cl, 150mM NaCl, 0.1% TWEEN 20) for 2 h, the membrane was incubated with an antibody against GFP (1:5,000 dilution) for 2 h at room temperature. The GFP antibody was already conjugated to the horseradish peroxidase (Anti-GFP-HRP, http://www.miltenyibiotec.com, #130-091-833). Blots were washed with TBST three times for 15 min each and developed with an enhanced chemiluminescence system according to the manufacturer (Bio-Rad, Hercules, CA).

### Quantification of RNA

Total plant RNA extraction and DNAse treatment were performed using the SV Total RNA Isolation Kit (Promega, Madison, WI, USA), followed by Reverse Transcription with SMART ® MMLV (Takara Bio USA, Mountain view, CA, USA). cDNA was synthesized using Oligo dT from 1 μg of total RNA. Viral RNA and GFP transcripts were quantified relative to the actin transcripts using reverse transcription quantitative real-time PCR (RT-qPCR). All the primers used for quantification are listed in Table S4. RT-qPCR was performed using the Bio-Rad CFX384™ Real-Time System in a 10-μL mixture containing SYBR Green PCR Master Mix (Applied Biosystems, Foster City, CA, USA). The thermocycling conditions were: 2 min polymerase activation at 50 °C followed by initial denaturation for 2 min at 95 °C and 45 cycles at 95 °C for 15 s, 60 / 55 °C for 1 min. Each sample was quantified in triplicates and a no template control was included. Cycle time values were automatically determined for all plates and genes using the Bio-Rad CFX384™ Real-Time System software. Analysis of RT-qPCR fluorescence data was performed and expressed in fold change relative to actin using the ΔΔCT method (Livak and Schmittgen 2001).

### Phytohormone Analysis

Phytohormones were extracted using a protocol mentioned in Bera *et al*., 2020. Briefly, frozen samples were homogenized with one milliliter of extraction buffer (isopropanol:water:HCl [2:1:0.005, v/v]) along with internal standards (d_4_-salicylic acid and d_5_-jasmonic acid; CDN Isotopes). After dichloromethane extraction, samples were redissolved in 100μL of methanol, and 10μL was analyzed using an Agilent Technologies 6420 triple quad liquid chromatography-tandem mass spectrometry instrument (Agilent, Santa Clara, CA, USA) as in Patton *et al*., (2019).

### RNA-seq experiment

Wild-type Arabidopsis (*Arabidopsis thaliana*) Columbia-0 were obtained from the Arabidopsis Biological Resource Center (http://www.arabidopsis.org). After 3 weeks of growth, one-half of the plants was infected with TuMV-GFP as described above. After 1 week, infected plants were identified by fluorescence under UV light. For aphid induction, 15 adult apterous aphids were caged on one leaf per plant for six different plants (uninfected plants). A corresponding set of six plants received cages with no aphids as a mock-inoculated control and six TuMV-GFP infected plants received cages to control for cages on the other plants. Caged leaves were developmentally matched, and infected leaves were verified for full infection before caging based on GFP visualization. Forty-eight hours after aphid placement, cages and aphids were removed and two leaves were pooled for each sample resulting in 3 replicates of pooled leaves for each treatment. RNA was then extracted as described above.

### Library Preparation, and Sequencing

Sequencing libraries were prepared using a multiplexing library protocol (Zhong *et al*., 2011). Briefly, oligo(dT) Dynabeads were used to purify mRNA, which was then fragmented, and the first-strand cDNA was synthesized using random primers, dNTP, and reverse transcriptase. The second-strand was synthesized using a dUTP mix, DNA Polymerase I, and RNase H, ends repaired, and adenylated. The cDNA fragments were ligated to adapters, selectively enriched by PCR, and purified using 138the AMPure XP beads. The library quality was assessed using the Agilent Bioanalyzer 2100 system and sequenced using an Illumina HiSeq 2000 instrument

### RNA-Seq data analysis

The quality of the raw reads was assessed with FASTQC and ShortRead. All samples presented reads with high quality. Reads were mapped against *Arabidopsis thaliana* TAIR10 genome using TopHat2 (Kim *et al*., 2013). The number of reads per gene was counted using HT-Seq and normalized using the normalization method implemented inside the edgeR Bioconductor package. The clusterization profile of the normalized samples was verified by Principal Component Analysis (PCA) and Spearman correlation. Differential expression test was conducted using edgeR, according to (Anders *et al*., 2013), using mock-infected samples as the reference control treatment. Genes with a FDR-corrected p-value lower than 0.1 were considered as differentially expressed genes (DEGs). Reads are available at the NCBI SRA (PRJNA60524).

### Gene Set Enrichment Analysis (GSEA)

To identify molecular mechanisms potentially relevant to the plant response to TuMV and aphids, a GSEA was conducted. The GSEA identified biological processes (BPs), molecular functions (MFs) and cellular components (CCs) that were over-represented among a list of DEGs. Categories with a p-value lower than 0.005 in a hypergeometric test were considered enriched.

### Protease and protease inhibitor assays

Twenty *Nicotiana benthamiana* plants that were 4-weeks old were agro-infiltrated with TuMV as described earlier. Five days post infiltration, the third youngest leaf of ten plants each were agro-infiltrated with 6K1-GFP or empty vector containing GFP construct (EV-GFP) and 100mg of plant tissue were collected 60hrs post inoculation. To evaluate the effect of proteases in virus infected plants in early stages of infection, twenty four 4-week old plants were also co-infiltrated with TuMV and either GFP or 6K1 (twelve plants for each treatment), and tissues were collected after 60hr. Plant tissues were homogenized in 1ml of 0.046M Tris-HCl and 0.0115M CaCl_2_ buffer (pH=8.1) with 5% polyvinylpolypyrrolidone. The homogenized samples were incubated on ice for 10mins followed by centrifugation at 11,000g for 10mins at 4C. The supernatant containing the soluble proteins from the leaves were then used for assays. Total protein extracted in each sample was measured by Bradford assay (Bradford, 1975). Fifty microliters of the protein extract were used to measure total protease activity in each sample using FITC Casein according to manufacturer’s protocol (Sigma Aldrich, USA). Known concentrations of trypsin was used as standards for protease assay. Protease activity in each sample was reported as equivalent amount of trypsin activity per mg of total protein.

### Statistical Analysis

The distribution of all values for all variables was analyzed to test for normality using the Shapiro-Wilk test (Sokal and Rohlf 1995) and were also tested for homogeneity of variances using the Levene test (Sokal and Rohlf, 1995). To determine if 6K1 expression impacts virus infection in local and systemic leaves (Fig. 5), the data were analyzed by generalized linear models (GLM) with a normal distribution curve which fitted the observed data. The model included treatments (TuMV, GFP, 6K1:GFP) and leaves (local, systemic) as fixed factor in a full factorial model. The GLM analysis was selected because it is a robust method with respect to the distribution of the data and allows contrasting both balanced and non-balanced models. To determine if the observed differences between classes of the same factor were significant, least significant difference (LSD) analysis were performed. The data related to Fig. 1, 2, and 4 were analyzed either by *t*-test or Kruskal-Wallis test. To determine if the protease activity between 6K1-GFP and EV-GFP treated plants were different from each other (Fig. 4C,D,F), the data was log transformed to meet assumptions of normality and a one-way ANOVA was performed at α<0.05 using R. The statistical analyses were performed using the SPSS v.24.0 program (SPSS Inc., IL, USA).

## Acknowledgments

This work was supported by NSF award 1723926 to CLC. We thank Jacob Tracy for excellent technical assistance.

## Author contributions

CLC and SB conceived the project. SB and CLC designed the research. SB, GA, SF, SR, and CLC performed the research and analyzed the data. CLC and SB interpreted the data. CLC and SB wrote the article.

---

**Fig. S1.**
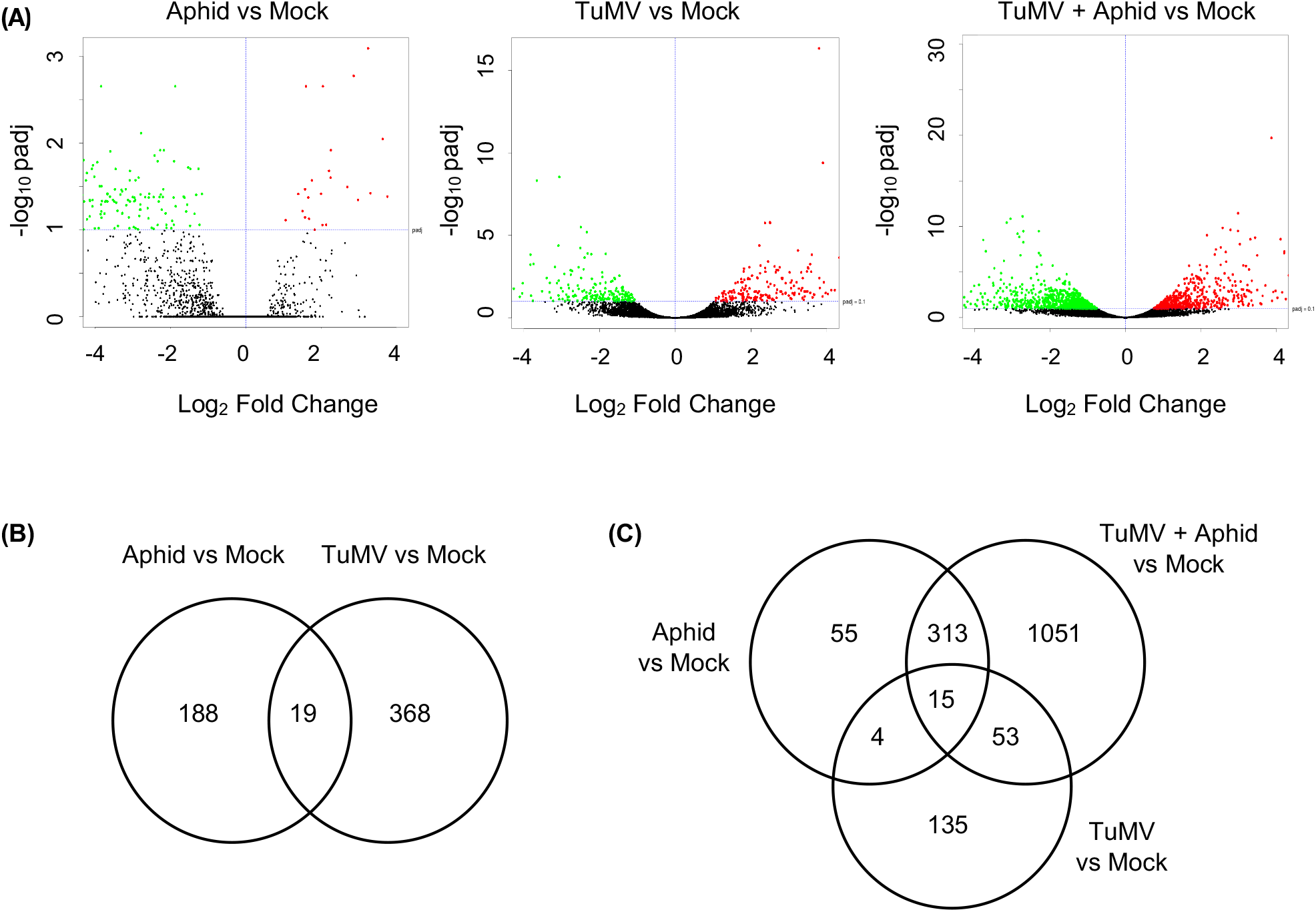
(A) Volcano-plots of −log_10_p and log_2_FC exhibited by each gene in *Arabidopsis thaliana* with aphids, TuMV, *o*r both aphids and TuMV, compared to mock controls. Up-and down-regulated genes are presented in red and green, respectively. (B) Numbers of differentially expressed genes (DEGs) shared between aphid-infested and TuMV-infected *A. thaliana* compared to mock controls. (C) Numbers of differentially expressed genes (DEGs) shared among all three treatments compared to mock controls. DEGs were identified using DESeq2and defined by |log2FC| ≥ 1; false discovery rate (FDR)-corrected p-value ≤ 0.1. FC, fold-change; p, FDR-corrected p-value.

**Fig. S4.**
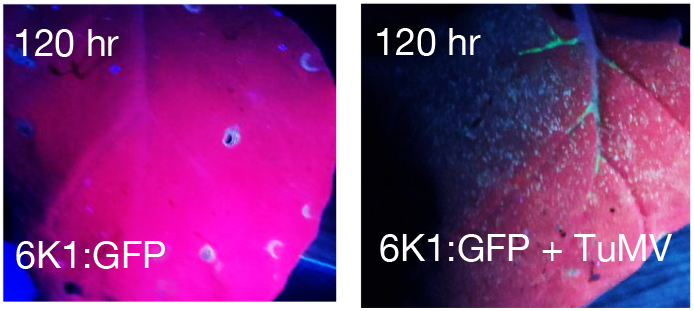
The construct 6K1:GFP was co-infiltrated with or without TuMV. Pictures were taken under a UV lamp of the agro-infiltrated leaves 120 hr later. The green fluorescence indicates the protein accumulation of 6K1:GFP.

**Table S4.**
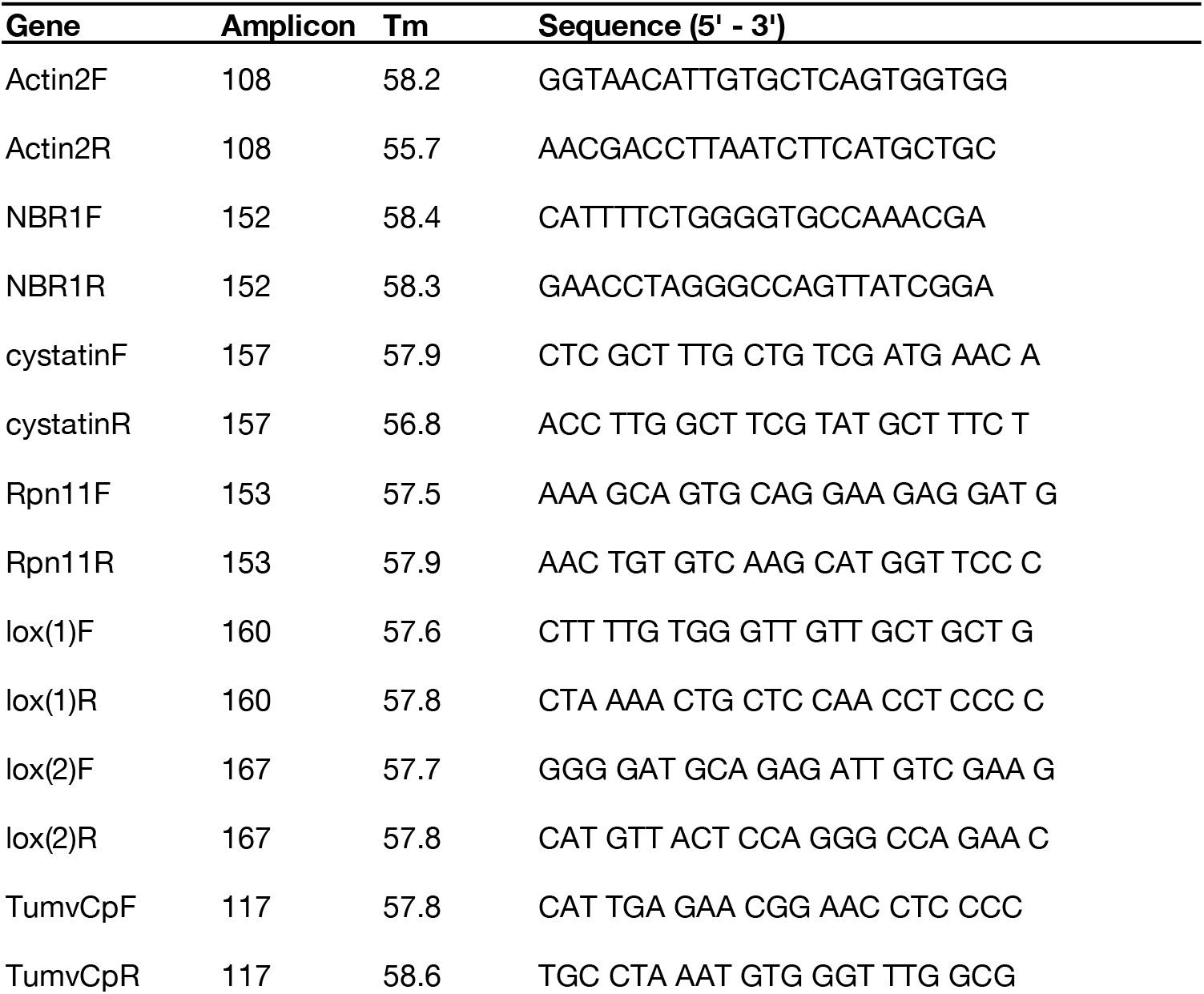
A list of primers used for quantification.

